# Double dissociation of fMRI activity in the caudate nucleus supports *de novo* motor skill learning

**DOI:** 10.1101/2019.12.24.887786

**Authors:** Yera Choi, Emily Yunha Shin, Sungshin Kim

## Abstract

Motor skill learning involves a complex process of generating novel movement patterns typically guided by evaluative feedback such as reward. Many studies have suggested that two separate circuits in the basal ganglia, rostral and caudal, are implicated in early goal-directed and later automatic stages of motor skill learning, respectively. However, there remains much to be elucidated about the respective involvement of the basal ganglia circuits in learning motor skills from scratch, which requires learning arbitrary action-outcome associations. To investigate this issue, we conducted a novel human fMRI experiment in which the participants learned to control a computer cursor on a screen by manipulating their right fingers. The experiment consisted of two fMRI sessions separated by five behavioral training sessions over multiple days. We discovered a double dissociation of fMRI activities in the rostral and caudal caudate nucleus, which were associated with skill performance in the early and late stages of learning. Moreover, we found that cognitive and sensorimotor cortico-caudate interactions distinctively predicted individual learning performance. In line with recent non-human primate studies, our results support the existence of parallel cortico-caudate networks involved in goal-directed and automatic stages of *de novo* motor skill learning.

## Introduction

Motor skill learning is a complex process of acquiring new patterns of movements to achieve more accurate and faster performance for motor tasks through repetitive practice^1^. It is an adaptive mechanism crucial for the survival and well-being of all animals, as it enables more efficient motor behaviors, which can lead to favorable outcomes such as greater rewards per unit time ^2^.

Existing literature has revealed that the dynamic interplay of multiple brain regions—encompassing the frontoparietal cortices, cerebellum, and basal ganglia—is required for the acquisition of new motor skills ^3,4^. The cortico-basal ganglia circuit is of particular interest, as many anatomical findings ^5,6^, animal studies ^7-10^, and human functional magnetic resonance imaging (fMRI) studies ^4,6,11-18^ have shown that distinct patterns of interaction between the structures in this circuit arise in different stages of learning. Specifically, evidence suggests that a transition from the rostral associative to the caudal sensorimotor (in rodents, from the dorsomedial to the dorsolateral) regions in the basal ganglia occurs during learning.

Notably, many studies that have reported the differential involvement of the separate cortico-basal ganglia circuits during motor skill learning have adopted sequence learning tasks ^11,12,14-16,18,19^, in which participants practiced repetitive sequences of well-learned discrete actions such as button-press. Although such sequence learning paradigms have elucidated many important aspects of motor learning, they may provide limited explanations for the initial acquisition of continuous motor skills involving explorative action selections ^1,20^.

Meanwhile, relatively few studies employed *de novo* motor learning paradigms, in which participants learned arbitrary associations between stimuli and required actions ^21-24^. Accordingly, there remains much to be elucidated about the neural basis of *de novo* motor learning. In this type of learning, action selection is sensitive to evaluative feedback such as reward, and thus the basal ganglia circuits processing reward are particularly important, as implied by studies in patients with motor-related neurodegenerative disorders ^25^. Consistent with the idea of functional dissociation in the basal ganglia for motor learning, recent non-human primate studies also found that the rostral and caudal regions of the caudate nucleus respectively modulate early flexible and later stable values arbitrarily associated with visual stimuli ^26,27^. The separate circuits mediate the transition from early goal-directed behavior to later automatic behavior, resulting in improvement of skill performance.

The present study thus aimed to investigate whether the human caudate nucleus serves distinct functions at different stages of continuous *de novo* motor skill learning. We conducted a series of behavioral and fMRI experiments, in which participants learned to control an on-screen cursor via an MR-compatible data-glove interface. Hand postures recorded by fourteen sensors of the data-glove were linearly mapped onto cursor locations. As a result, we demonstrated a clear double dissociation of success-related fMRI activities in the caudate nucleus, which declined in the rostral region but increased in the caudal region from the early to late stages of learning. Furthermore, we also found that the intrinsic functional interactions between the caudate nucleus and cortical regions were predictive of individual learning performance. In sum, our findings suggest that the caudate nucleus serves as the main locus for *de novo* motor skill learning in humans.

## Results

### Successful learning of de novo motor skills

Thirty participants completed the entire study, which included two fMRI sessions separated by five behavioral training sessions outside the scanner (Figure 1). In the first and second fMRI sessions, participants were exposed to two different mappings between the cursor and finger movements while performing the main task, for which they were asked to reach on-screen targets with the cursor by moving their right fingers. During the behavioral training sessions, participants were trained only for one of the two mappings (i.e., the practiced mapping), and also performed the sequential task in addition to the main task. See **Materials and Methods** for detailed information on the experimental design.

**Figure 1.**
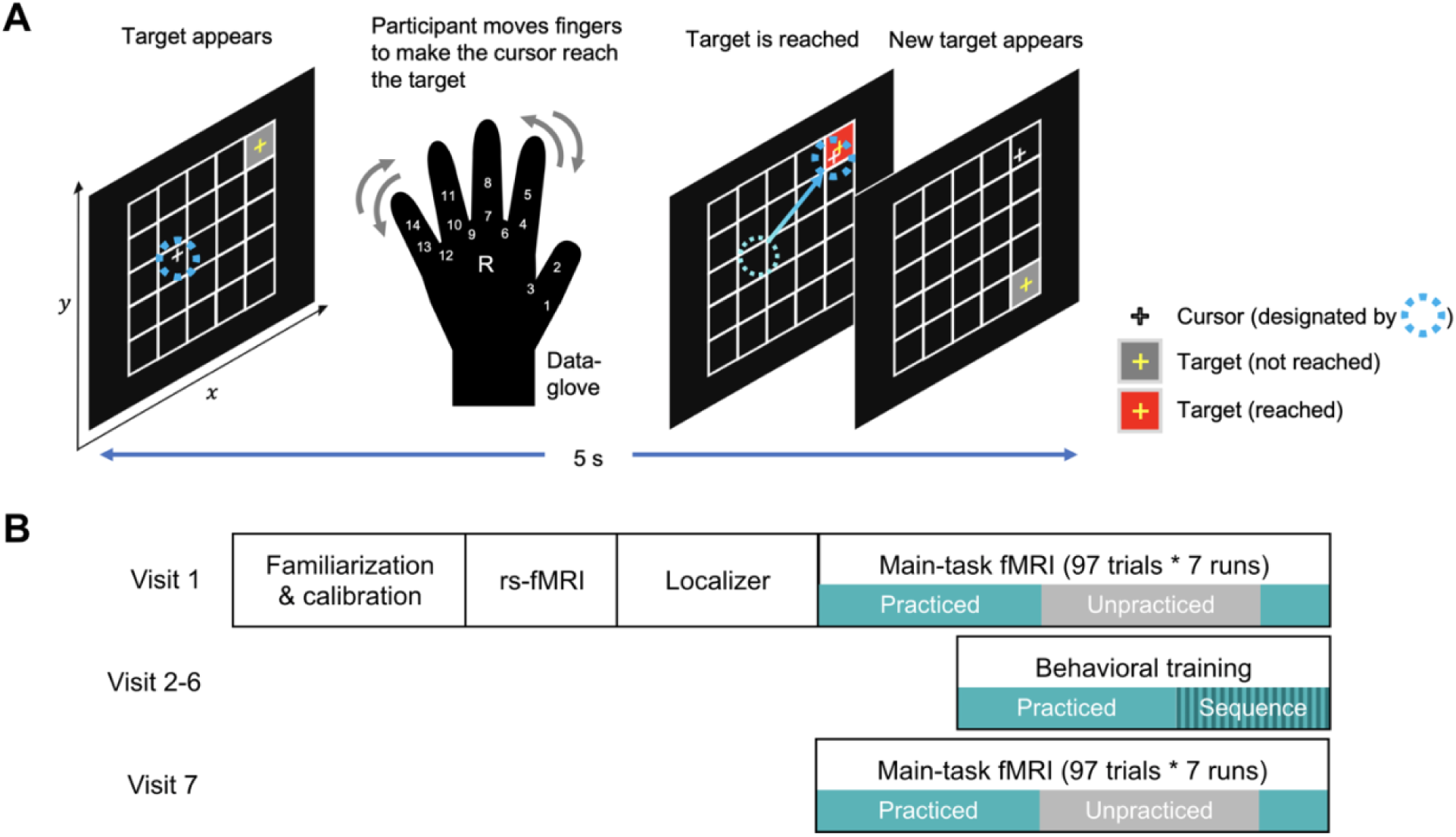
Overview of the experiment design. **A.** Overview of the *de novo* motor learning task. Participants learned to control a cursor (white crosshair) to reach a target (gray rectangular grid cell with a yellow crosshair at its center) on a 5 × 5 grid by manipulating their right fingers, using a data-glove measuring degrees of freedom with 14 sensors on finger flexures and abductions. The 14-dimensional vector representing hand posture was linearly mapped onto the 2-dimensional position of the cursor. When the cursor was on a target grid cell, the color of the target cell was changed to red. Participants were instructed to stay on the target as long as possible and try to get as close as possible to the center of the target. **B**. Visit schedule of the entire experiment. **Visit 1 (fMRI 1)**: After the familiarization and calibration phases to determine and inspect the cursor-hand posture mapping created from random finger movements, an fMRI session composed of a resting-state fMRI scan, a localizer scan to identify regions implicated in random finger movements, and seven runs of the main-task fMRI scan including both of the practiced (green) and unpracticed (gray) mappings followed. **Visits 2-6 (Behav 1-5)**: A behavioral training session only on the “practiced” mapping (including three runs of the main task and two runs of the sequential task) was conducted outside the MRI scanner. **Visit 7 (fMRI 2)**: Seven runs of the main-task fMRI, including both “practiced” and “unpracticed” mappings, were conducted.

Actual cursor trajectories made by a representative participant demonstrated evident learning of the ability to maneuver the cursor to the targets, which significantly improved from the first fMRI session to the intermediate behavioral and the second fMRI sessions (Figure 2A). All individual participants demonstrated significant improvement of performance in terms of the straightness of cursor trajectories (Figure 2B) and the success rate (Figures 2C and S1), primarily for the practiced mapping. The success rate, which was calculated as the fraction of time when the cursor stayed on the targets and used as the major indicator of task performance, tended to increase across the entire experimental sessions. This increase suggests that the participants successfully acquired the ability to reach the target grid cells quickly and stably for the practiced mapping. Between-session forgetting, which was observed as a drop of task performance at the beginning of each session, tended to decrease across sessions, as shown in our previous study using motor adaptation tasks ^28^.

**Figure 2.**
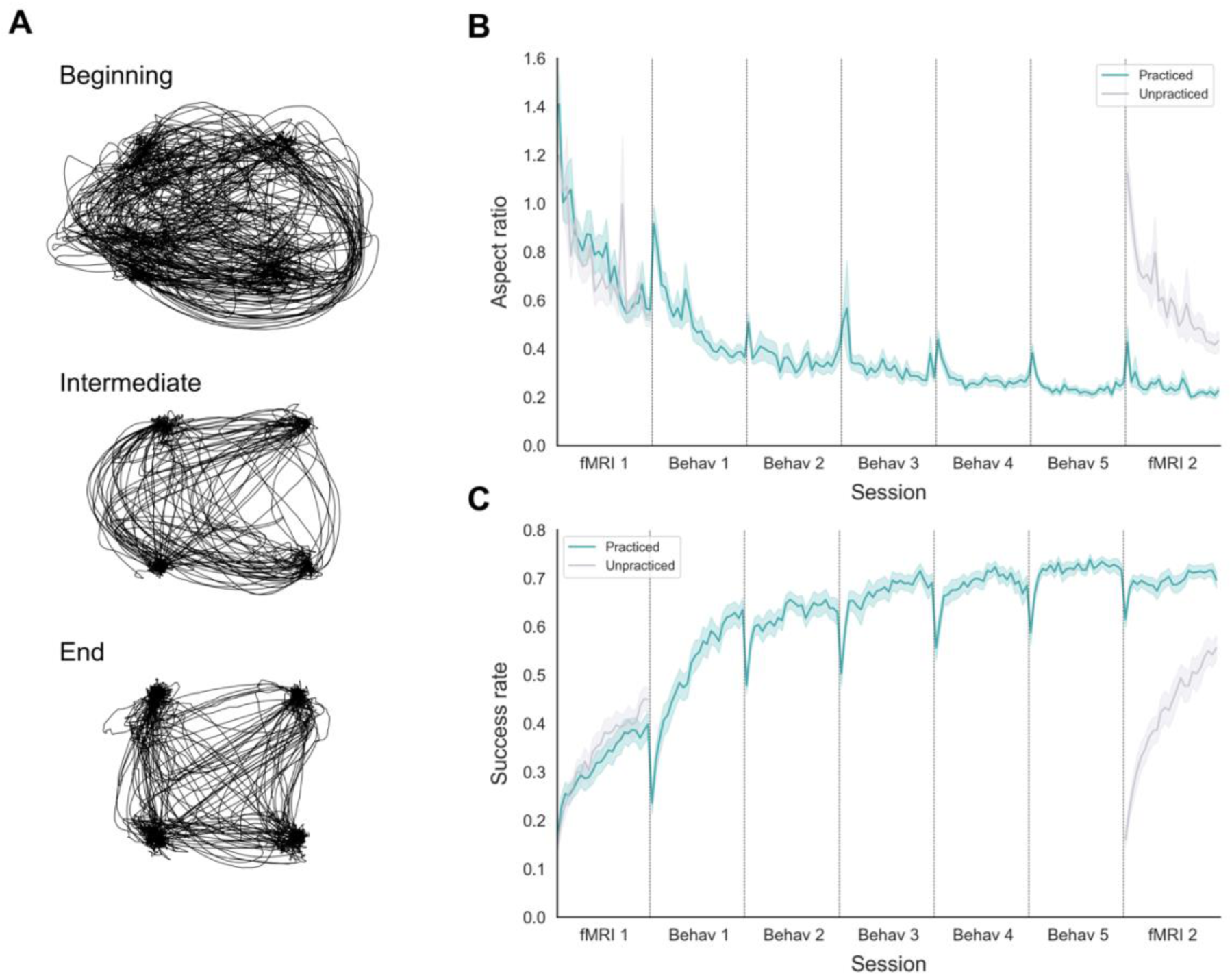
Behavioral results. **A.** Sample of actual cursor trajectories for the practiced mapping made by one representative participant, at different learning stages (Beginning: fMRI 1, Run 1; Intermediate: Behav 3, Run 2; End: fMRI 2, Run 3). **B**. Curve of the block-by-block aspect ratio (i.e., the maximum perpendicular distance between the straight line connecting two consecutive targets and the actual cursor trajectory normalized by the length of the straight line), a measure of straightness, for different mappings. The green solid line denotes the group average of the block-by-block aspect ratio for the practiced mapping, while the gray solid line denotes that for the unpracticed mapping. Lighter-colored shades around the solid lines indicate SE. Since participants were only trained on the practiced mapping during the five behavioral sessions (Behav 1-5), only the data from the two fMRI sessions (fMRI 1, 2) exist for the unpracticed mapping. The decrease in the aspect ratio suggests that participants tended to make straighter movements as learning proceeded. **C**. Curve of the block-by-block success rate (proportion of time during which the cursor was on the target) for different mappings. The green solid line denotes the group average of the success rate for the practiced mapping, while the gray solid line denotes that for the unpracticed mapping. Lighter-colored shades around the solid lines indicate SE. The increase in the success rate suggests that participants tended to reach the targets more quickly and stayed longer on them as learning proceeded.

For the unpracticed mapping, to which participants were exposed only during the fMRI sessions, the learning curves showed patterns similar to those of the first two sessions of the practiced mapping. Indeed, there were no significant differences in the amount of learning between the two mappings in the first fMRI session (i.e., the first sessions of the mappings; *p* = 0.199, Wilcoxon signed-rank test), as well as between the unpracticed mapping in the second fMRI session and the practiced mapping in the first behavioral training session (i.e., the second sessions of the mappings; *p* = 0.797, Wilcoxon signed-rank test). Yet a significant between-mapping transfer of learning was observed, as the final amount of transfer was significantly greater than the initial amount of transfer (*p* = 0.003, Wilcoxon signed-rank test; see **Materials and Methods** for details).

### Spatiotemporal dissociation of the fMRI activity related to motor skill learning in the caudate nucleus

Seeking the goal of maximizing the success rate, the participants learned to select and execute a series of continuous actions leading to higher success rates, through the trial-and-error exploration of the action-outcome mapping. In this aspect, the success rates associated with different actions can be considered as “action values.” We thus used a parametric regressor modulating the time-varying success rate for the whole-brain voxel-wise general linear model (GLM) analyses.

The analyses identified distinctive patterns of fMRI activity in the early versus late stages of learning (Figures 3 and S2, Table 1). The regions showing significant success-rate modulation during early learning were comparable to those reported by previous studies ^29,30^: the cognitive-attentional network, including the bilateral supramarginal gyrus, middle cingulate cortex, insula, and superior parietal cortex, as well as subcortical regions including rostral putamen, rostral caudate nucleus, right thalamus, and other regions responsive to visual stimuli.

**Table 1.** Whole-brain voxel-wise GLM analysis comparing the strength of positive success-rate modulation between early versus late stages of learning.

|  | Peak (MNI) |  |  | Cluster size | Z-score | Corrected |
| --- | --- | --- | --- | --- | --- | --- |
|  | X | Y | Z | (voxels) | at peak | p-value |
| Early > Late |  |  |  |  |  |  |
| R Supramarginal Gyrus | 63 | -26 | 41 | 1059 | 6.17 | p < 0.01 |
| R Middle Cingulate Cortex | 9 | 25 | 33 | 878 | 5.33 | p < 0.01 |
|  | 12 | -31 | 41 | 102 | 4.96 | p < 0.01 |
| R Insula Lobe | 36 | 17 | 1 | 818 | 6.09 | p < 0.01 |
| L Supramarginal Gyrus | -66 | -26 | 38 | 793 | 5.84 | p < 0.01 |
| L Insula Lobe | -36 | 20 | -5 | 469 | 5.80 | p < 0.01 |
| R Putamen and Caudate | 17 | 9 | -7 | 284 | 6.29 | p < 0.01 |
| L Putamen and Caudate | -9 | 9 | -8 | 211 | 5.21 | p < 0.01 |
| R Inferior Temporal Gyrus | 47 | -56 | -16 | 147 | 4.70 | p < 0.01 |
| L Middle Temporal Gyrus | -50 | -58 | -2 | 116 | 4.73 | p < 0.01 |
| R Middle Occipital Gyrus | 44 | 82 | 25 | 102 | 5.03 | p < 0.01 |
| L Superior Parietal Lobule | -17 | -74 | 54 | 95 | 4.53 | p < 0.01 |
| L Inferior Occipital Gyrus | -31 | -99 | -10 | 63 | 5.23 | p < 0.02 |
| R Thalamus | 7 | -23 | 1 | 53 | 4.56 | p < 0.03 |
| R Angular Gyrus | 28 | -61 | 43 | 44 | 3.85 | p < 0.04 |
| R Inferior Occipital Gyrus | 36 | -88 | -5 | 43 | 4.33 | p < 0.05 |
| R Middle Frontal Gyrus | 34 | 49 | 33 | 43 | 4.27 | p < 0.05 |
|  | 47 | 46 | 6 | 41 | 4.18 | p < 0.05 |
| R Superior Parietal Lobule | 17 | -72 | 54 | 40 | 3.86 | p < 0.05 |
| Early < Late |  |  |  |  |  |  |
| R Caudate | 20 | -40 | 19 | 254 | 5.93 | p < 0.01 |
| L Caudate | -26 | -50 | 14 | 97 | 4.64 | p < 0.01 |
| L Thalamus | -4 | -29 | 11 | 43 | 4.25 | p < 0.05 |
| Common |  |  |  |  |  |  |
| Ventromedial cortex | -4 | 49 | 3 | 138 | 4.73 | p < 0.01 |
| L Inferior Parietal Lobule | -50 | -74 | 41 | 124 | 5.26 | p < 0.01 |
| L Superior Frontal Gyrus | -9 | 54 | 49 | 108 | 4.65 | p < 0.01 |
| R Inferior Parietal Lobule | 58 | -58 | 35 | 55 | 4.59 | p < 0.05 |
| R Superior Medial Gyrus | 4 | 52 | 52 | 50 | 4.25 | p < 0.05 |

Clusters showing significant positive modulatory effects of the success rate in the early (Early > Late) and later (Early < Late) stages of learning are listed (voxel-wise $p < 0.001$ , 40 suprathreshold voxels for cluster-corrected $p < 0.05$ ). In addition, to identify the regions significantly modulated in both stages of learning, a conjunction analysis was performed with $p < 0.001$ , and the average peak Z-scores across learning stages are presented for these regions.
Abbreviations: L, left; MNI, Montreal Neurological Institute; R, right.

**Figure 3.**
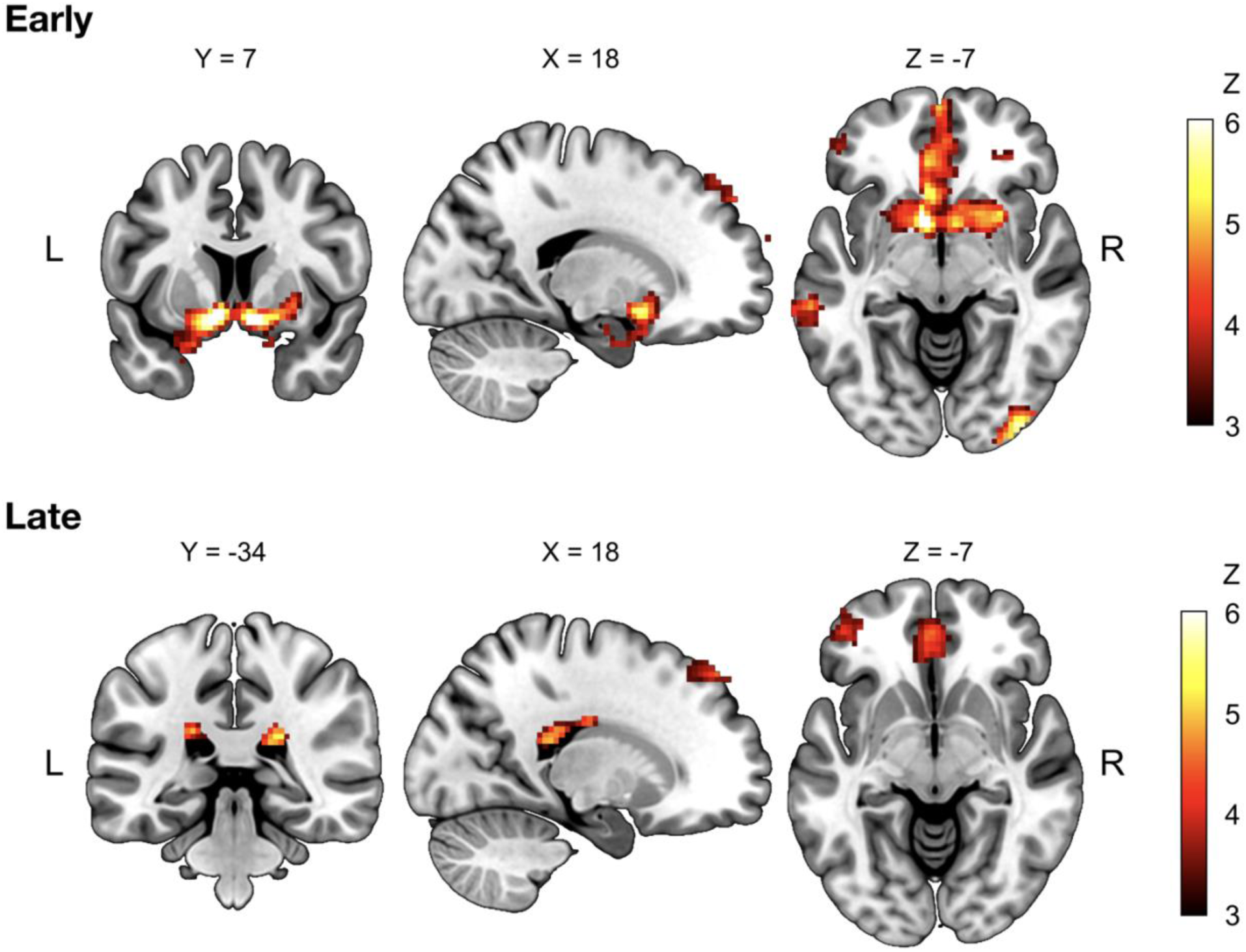
Whole-brain voxel-wise GLM analyses using a parametric regressor modulating success rates. Results from the whole-brain voxel-wise GLM analyses with a parametric regressor modulating the time-varying success rate estimated every 1 s (voxel-wise *p* < 0.001, cluster-corrected *p* < 0.05). Regions showing significant positive modulation by the success rate in the early (fMRI 1) and late (fMRI 2) stages of learning are presented. The success rate, which can be considered as an action value, was highly modulated by the activities in the rostral striatal regions including the caudate head and nucleus accumbens in the early stage, and by those in the caudate tail in the late stage. Interestingly, there were significant success-modulated activities in both stages of learning in the VMPFC, which is well-known for value- and reward-processing. Color bars indicate the group-level *Z*-scores, and numbers above each slice indicate MNI coordinates. Abbreviations: GLM, general linear model; L, left; MNI, Montreal Neurological Institute; R, right; VMPFC, ventromedial prefrontal cortex.

The fMRI activities related to the success rate globally decreased in the rostral striatal and prefrontal-parietal cortical regions as learning proceeded, potentially reflecting habituation and neural efficiency ^3,12,18,31^. Meanwhile, the modulatory activity locally increased in the bilateral caudal regions of the caudate nucleus (Figure 4, Table 1). Notably, the caudate nucleus demonstrated a gradual transition of the activity from the rostral to caudal regions (Figure 4). This finding is in line with previous literature on motor sequence learning that suggests early involvement of the anterior regions and later involvement of the posterior regions of the basal ganglia, including the putamen ^11^.

**Figure 4.**
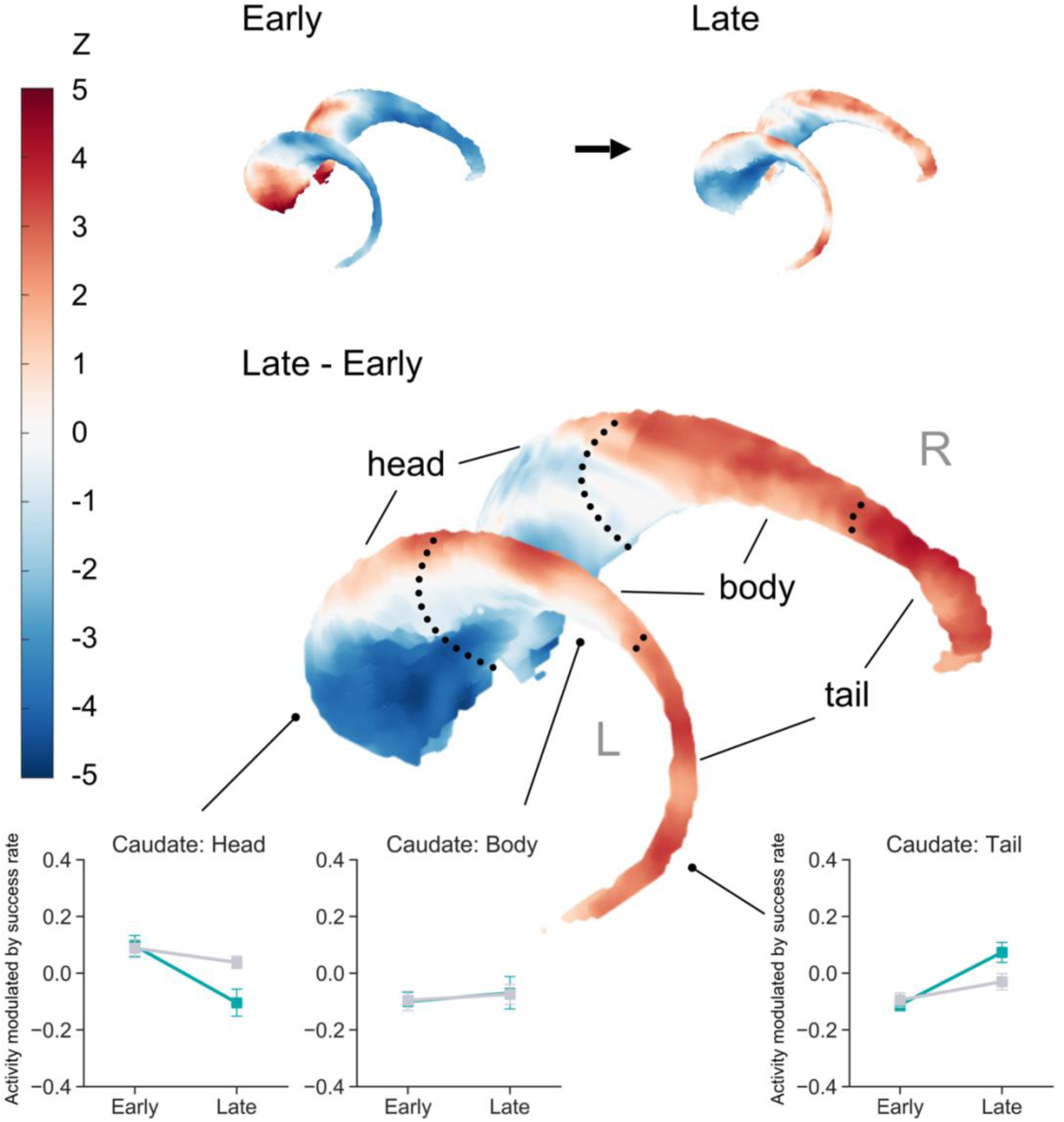
Spatiotemporal dissociation of the fMRI activity in the caudate nucleus. Rendering of the caudate nucleus illustrating the gradual transition of subregions modulating the success rate, from the rostral (head) to caudal (tail) regions. Upper panel: The left image (“Early”) represents the pattern of success-rate modulation in the early stage of learning, while the right image (“Late”) represents the pattern of success-rate modulation in the later stage of learning. Middle panel: Red and blue colors respectively indicate regions showing increased and decreased success-rate modulation from the early to late stage. The color bar indicates the group-level *Z*-scores converted from a paired *t*-test. Bottom panel: Activity modulated by the time-varying success rate in the caudate subregions, for the practiced (green solid line) and unpracticed (gray solid line) mappings in the early and later stages of learning. Error bars indicate SE. Abbreviations: L, left; R, right; SE, standard error.

To further examine the learning-induced changes in fMRI activity, we conducted region-of-interest (ROI) analyses, primarily in the subregions of the caudate nucleus as well as other regions frequently implicated in reward processing, such as the putamen, globus pallidus, nucleus accumbens (NAc), and ventromedial prefrontal cortex (VMPFC). For the detailed procedure of generating the ROIs, see **Materials and Methods**. As there exists evidence of the learning-induced transition of activity from the rostrodorsal to caudoventral regions of the putamen ^11^, subdivided ROIs of the putamen were defined accordingly.

Two-way (learning stage X mapping) repeated-measures ANOVA was conducted for each ROI to examine the effects of learning stage and mapping. In the caudate head, the main effects of learning stage (*F*(1,29) = 19.99, *p* < 10^−3^, η^p2^ = 0.41) and mapping (*F*(1,29) = 7.42, *p* = 0.011, η^p2^ = 0.20) were significant, yet their interaction was not significant (*F*(1,29) = 3.22, *p* = 0.08). Post-hoc tests revealed that the modulatory activity in the caudate head was significantly lower for the practiced mapping than for the unpracticed mapping in the later stage of learning (*T*(29) = 2.60, Holm-Bonferroni-adjusted *p* = 0.029, Cohen’s *d* = 0.76). Meanwhile, the caudate tail demonstrated a significant main effect of learning stage (*F*(1,29) = 24.25, *p* < 10^−4^, η^p2^ = 0.46) as well as a significant interaction between learning stage and mapping (*F*(1,29) = 4.95, *p* = 0.034, η^p2^ = 0.15), yet no significant between-mapping differences in the degree of modulation were found in post-hoc tests. The anterior putamen and globus pallidus also showed similar patterns, with significant main effects of learning stage (anterior putamen: *F*(1,29) = 22.32, *p* < 10^−4^, η^p2^ = 0.43; globus pallidus: *F*(1,29) = 16.10, *p* < 10^−3^, η^p2^ = 0.36) and interactions (anterior putamen: *F*(1,29) = 5.72, *p* = 0.023, η^p2^ = 0.16; globus pallidus: *F*(1,29) = 7.02, *p* = 0.013, η^p2^ = 0.19). In the case of the globus pallidus, the degree of modulation for the practiced mapping was significantly lower than that for the unpracticed mapping in the later stage (*T*(29) = 2.41, Holm-Bonferroni-adjusted *p* = 0.045, Cohen’s *d* = 0.66).

Only the main effect of learning stage (*F*(1,29) = 5.29, *p* = 0.029, η^p2^ = 0.15) was significant in the posterior putamen, yet post-hoc tests revealed no significant between-mapping differences in modulation. In the NAc, all the main (learning stage: *F*(1,29) = 28.36, *p* < 10^−4^, η^p2^ = 0.49; mapping: *F*(1,29) = 24.25, *p* = 0.002, η^p2^ = 0.28) and interaction (*F*(1,29) = 24.25, *p* = 0.002, η^p2^ = 0.29) effects were significant, and the late-stage modulatory activity was lower for the practiced mapping than the unpracticed mapping (*T*(29) = 4.40, Holm-Bonferroni-adjusted *p* < 10^−3^, Cohen’s *d* = 0.94). Neither significant main (learning stage: *F*(1,29) = 0.48, *p* = 0.494; mapping: *F*(1,29) < 10^−5^, *p* = 0.998) nor interaction (*F*(1,29) = 0.008, *p* = 0.930) effects were observed in the caudate body. The main (learning stage: *F*(1,29) = 0.11, *p* = 0.744; mapping: *F*(1,29) = 0.09, *p* = 0.769) and interaction (*F*(1,29) = 0.15, *p* = 0.705) effects for the VMPFC were also not significant.

As suggested by these results, the activity in the caudate head was positively modulated by the success rate in the early stage. Then, after extensive training, this modulatory effect decreased and even changed its direction. In contrast, the caudate tail demonstrated significant negative modulation by the success rate in the early stage, which changed to be positive in the later stage. Indeed, additional two-way (ROI X learning stage) repeated-measures ANOVA contrasting only the caudate head and tail data for the practiced mapping revealed no significant main effects (ROI: *F*(1,29) = 0.14, *p* = 0.715; learning stage: *F*(1,29) = 0.04, *p* = 0.840), yet the interaction was highly significant (*F*(1,29) = 38.97, *p* < 10^−6^, η^p2^ = 0.57). The following post-hoc tests further suggested that this interaction was due to the early-stage positive modulation in the caudate head (*T*(29) = 3.56, Holm-Bonferroni-adjusted *p* = 0.001, Cohen’s *d* = 0.87) as well as the early-stage negative modulation in the caudate tail (*T*(29) = 4.43, Holm-Bonferroni-adjusted *p* < 10^−3^, Cohen’s *d* = 1.03).

Meanwhile, the success-related activity in the anterior putamen showed a significant decrease through the course of learning. Notably, however, in the posterior putamen, we did not find any significant change providing supporting evidence of the rostrodorsal-caudoventral transition in the putamen as suggested by previous literature on motor sequence learning. In addition, interestingly, the VMPFC, which is known to play a crucial role in value processing, did not show learning-induced changes in its success-related activity, which was significantly positive in both early and late stages of learning as demonstrated by a conjunction analysis (for the entire regions identified by the analysis, see the “Common” section of Table 1).

### Cortico-caudate functional interactions predict individual learning performance

Following our observation of the robust learning-induced rostrocaudal transition of fMRI activity in the caudate nucleus, we sought to address whether the respective interactions within the two separate cortico-caudate loops—the rostral cognitive and caudal sensorimotor loops—can predict individual learning performance (Figure 5A). For this, we used independently defined cortical ROIs, including the bilateral dorsolateral prefrontal cortex (DLPFC) and left motor/somatosensory cortex (M1/S1) (Figure S3), and performed functional connectivity analyses with resting-state fMRI data acquired prior to the first main-task fMRI session (Figure 1B).

**Figure 5.**
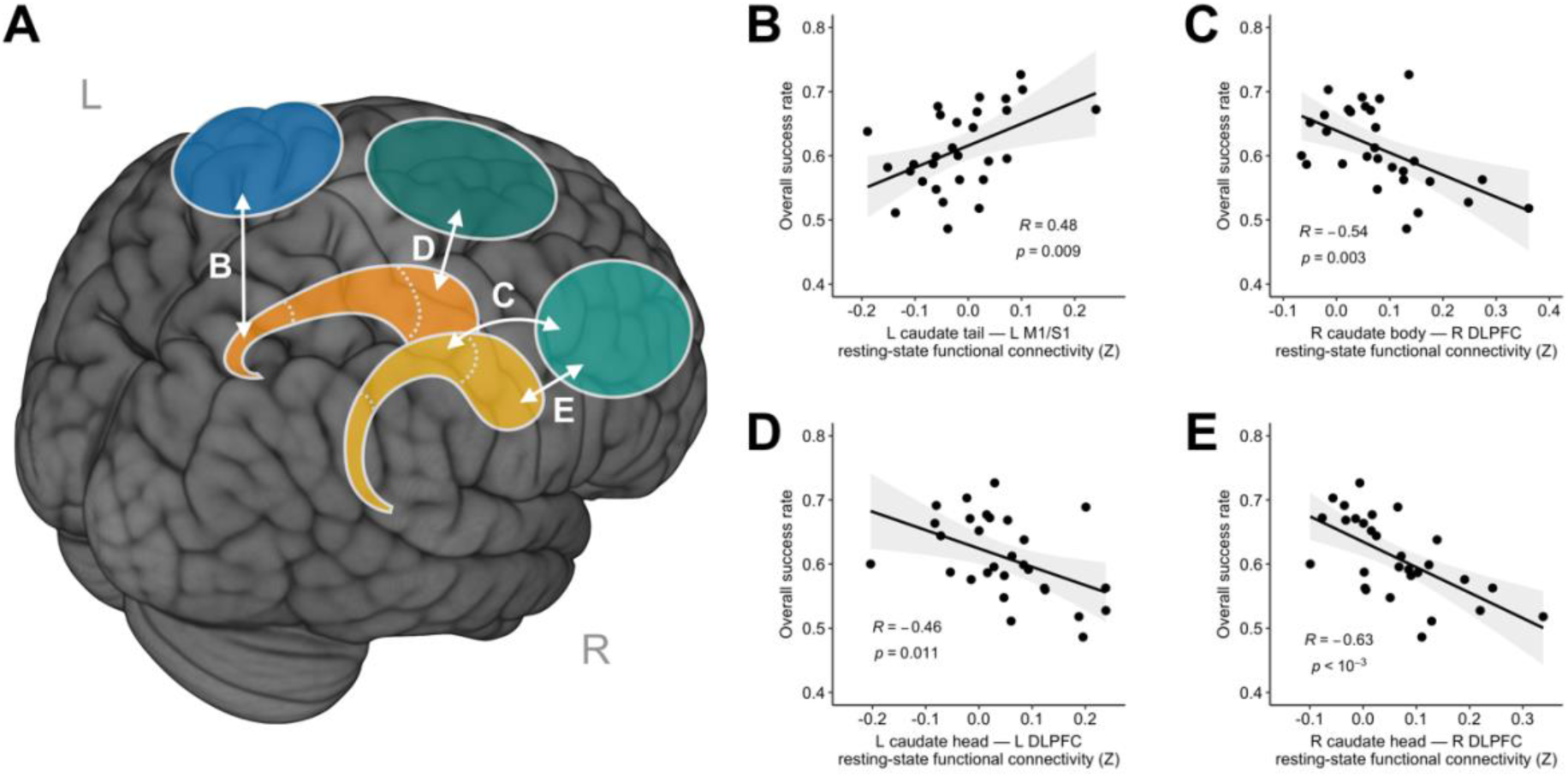
rs-fMRI connectivity predictive of individual learning performance. The relationship between the cortico-caudate intrinsic functional connectivity and learning performance. **A**. A schematic of seed ROIs in the caudate nucleus and the independently defined cortical ROIs in the left/right DLPFC and left M1/S1. **B**. The relationship between the L caudate tail — L M1/S1 rs-fMRI connectivity and the overall success. **C**. The relationship between the R caudate body — R DLPFC rs-fMRI connectivity and the overall success rate. **D**. The relationship between the L caudate head — L DLPFC rs-fMRI connectivity and the overall success rate. **E**. The relationship between the R caudate head — R DLPFC rs-fMRI connectivity and the overall success rate. The gray shades indicate 95% confidence interval. Uncorrected *p*-values are presented. Abbreviations: DLPFC, dorsolateral prefrontal cortex; L, left; M1, primary motor cortex; rs-fMRI, resting-state functional magnetic resonance imaging; R, right; S1, primary somatosensory cortex.

The intrinsic functional connectivity between the left caudate tail and the left M1/S1 showed significant positive correlations with the overall success rate (Figure 5B, *R* = 0.48, *p* = 0.009). Interestingly, however, the ipsilateral connectivities between the rostral regions of the caudate nucleus and the DLPFC showed significant negative correlations with the overall success rate (right caudate body, Figure 5C: *R* = −0.54, uncorrected *p* = 0.003, Bonferroni-corrected *p* = 0.01; left caudate head, Figure 5D: *R* = −0.46, uncorrected *p* = 0.011, Bonferroni-corrected *p* = 0.044; right caudate head, Figure 5E: *R* = −0.63, uncorrected *p* < 10^−3^, Bonferroni-corrected *p* = 0.001). The connectivity between the right caudate body and the right DLPFC also demonstrated a significant negative correlation with the overall success rate (Figure 5C: *R* = −0.54, uncorrected *p* = 0.003, Bonferroni-corrected *p* = 0.025). Taken together, these results further support parallel but dissociable processing of the cognitive and sensorimotor loops in the caudate nucleus in continuous *de novo* motor skill learning.

### Localization of hand movement-related regions

Using the data from an independent localizer scan (described in detail in **Materials and Methods**), regions significantly activated by random finger movements were identified by a group-level one-sample *t*-test (performed by AFNI’s *3dttest++* function with “-Clustsim” option), at a highly stringent voxel-wise threshold of *p* < 10^−5^ and a cluster size larger than 150 voxels. As a result, seven clusters were defined in the bilateral pre/postcentral gyrus, left posterior putamen, right cerebellum (lobules IV, V, and VIII), SMA, and thalamus (Table S1, Figure S3B). All of these regions are implicated in hand movement and have been shown to exhibit impairments in activation and connectivity under conditions such as focal hand dystonia ^32^.

In our experiment, it is notable that participants were required to stop moving their fingers once they reached a target until the next target appeared to obtain higher success rates. Due to this experiment design, the amount of movement was highly collinear with the success rate used for the GLM analysis (*R* = −0.51±0.02, mean±SE), indicating a greater amount of movement in the early stage of learning. Thus, we analyzed the localizer data to examine whether finger movements were sufficient to induce the dissociative activities in the caudate nucleus observed in the preceding analyses. Patterns of activity similar to those of success-related activities were observed neither in the caudate head (i.e., greater activity with increased finger movements in the early stage) nor in the caudate tail (i.e., greater activity with paused or reduced finger movements in the late stage) (*p* > 0.5 for both, one-tailed *t*-test). This result demonstrates that the double dissociation of fMRI activities in the caudate nucleus is less likely due to movement *per se*, but rather due to the learned values of motor actions as we hypothesized.

## Discussion

The current study investigated the role of the human caudate nucleus in *de novo* motor skill learning. The motor skill learning paradigm using the data-glove interface, which was adopted for the first time in an fMRI study, allowed participants to learn completely new mappings between their actions and outcomes as they pursued positive feedbacks in a continuous task space. Hence, the current study offers a unique opportunity to tract the neural changes during the emergence and development of a continuous motor skill, which may more closely resemble the complexity and diversity of motor skills in the real world than other tasks frequently implemented in laboratory settings. We discovered a robust learning-induced double dissociation of fMRI activities in the subregions of the caudate nucleus, as the success-rate modulation increased in the caudal region but decreased in the rostral region. Moreover, the interactions between these caudate subregions and cortical regions distinctively predicted individual learning performance, in consistent with their respective roles in the early goal-directed and late automatic stages of learning. Our results are the first demonstration of the role of the caudate nucleus in human *de novo* motor skill learning, which is in line with many human fMRI and non-human primate studies suggesting the parallel operation of the rostral cognitive and caudal sensorimotor loops in the basal ganglia in the early and late stages of sequence learning ^4,7,11,15-18,33,34^.

The absence of significant effects for the unpracticed mapping appears to preclude the possibility that the current findings are simply due to prolonged exposure to the task. However, we cannot completely rule out the possibility that the reduced amount of movement, rather than learning, might have exerted a considerable influence on the results. Nevertheless, null findings from the localizer task—which was performed without goal-directed movement—strongly suggest that the observed fMRI activities are not solely due to movement *per se*. Instead, these activities are more likely due to learning of goal-directed movement while maximizing utility, or reward per effort and time. Future studies with more deliberate experiment designs would be necessary to dissociate out movement *per se* from its learned utility and delineate the pure learning-induced activity in the caudate nucleus.

Movement vigor, which is suggested to be determined by movement-associated reward and effort ^35,36^, also accounts for this complex relationship between movement and utility in motor learning; movement vigor would increase either if reward increases or required effort decreases. In our experiment, to increase the success rate while reducing inefficient movement (i.e., “learning to be lazy”) would yield better skill performance. As the basal ganglia have been continuously implicated in the control of movement vigor ^37^, the rostrocaudal shift observed in the basal ganglia during learning may play an important role in maintaining an adequate level of movement vigor for desirable movement regardless of learning stage. Notably, there have been reports demonstrating reduced movement vigor in Parkinson’s disease (PD), which is characterized by bradykinesia and impairments in dopaminergic projections to the striatum ^38^.

Recently, in a series of primate studies, distinct sets of dopaminergic neurons innervating the rostral and caudal regions of the caudate nucleus were identified as the potential neurobiological underpinnings of the respective modulation of early flexible and later stable values for the learning of arbitrary object-value associations, which is named “object skill” ^26,27,39^. In these studies, monkeys learned to make saccades to multiple visual objects with higher chances of receiving liquid reward. The object-value contingency was consistent in the stable condition but was frequently reversed in the flexible condition. The neurons in the caudate head were found to be sensitive to immediate reward outcomes in the flexible condition, responding more strongly to high-value objects. Interestingly, after extensive training in the stable condition, the neurons in the caudate tail responded to the stably high-valued objects even in the absence of reward, as monkeys continued to show automatic gaze to the objects. These findings have significantly advanced the understanding of the role of the parallel basal ganglia circuits in mediating goal-directed and automatic processes for object skill. However, there remains much to be elucidated about the respective roles of these circuits for the “action skill” (i.e., the learning of arbitrary action-outcome associations). Here, we have demonstrated that successful learning of actions arbitrarily associated with higher values (i.e., greater success rates) was closely linked to the activities encoding “action values” in the caudate nucleus, which could be similarly dissociated for the early-flexible and late-stable stages of learning. These results provide the first evidence in humans supporting the hypothesis that the object and action skills share common neurobiological mechanisms involving the parallel circuits of the caudate nucleus.

Our results may also be interpreted in relation to the role of the caudate nucleus in well-studied category learning, since the current task involves feedback-based learning of associations between different hand postures and distinct target positions. It has been shown that impairments of the caudate head, which are typically observed in patients with PD, may lead to deficits in rule-based and explicit category learning ^40,41^. Few fMRI studies also have demonstrated that the activities in the body and tail of the caudate are associated with improved category learning ^42,43^. Yet, it should also be noted that, in our experiment, participants learned the general mapping between hand postures and cursor positions, rather than simple associations between specific hand postures and target positions, as suggested by the learning-induced increase in the straightness of movement (Figure 2A, B). Moreover, evidence of generalization to untrained targets was also observed in an additional analysis using the data from the main tasks as well as from the sequential tasks, in which adjacent untrained targets had to be reached sequentially along with the main-task targets (see **Materials and Methods** for details). This ability of generalization is a hallmark of implicitly acquired motor learning, which involves sensorimotor and corticostriatal networks, and has been well demonstrated by previous studies ^23,44^.

We hypothesized that the functional connectivity between the caudate head/body and prefrontal cortices would play an important role in implementing cognitive strategies in the early stage of learning, while the connectivity between the caudate tail and sensorimotor regions would be implicated in the later stage with more efficient and automatized movement. If an individual continues to rely heavily on conscious cognitive processes even in the later stage of learning, automatization of desired motor behavior may be delayed or hindered, which would in turn negatively affect performance ^45-47^. Conversely, if attention is divided by dual tasks and cannot be solely allocated to cognitive processes, automatic skill performance may improve ^48^. In line with these predictions, we found that participants with higher learning performance would show weaker intrinsic connectivity between the caudate head/body and DLPFC and stronger connectivity between the caudate tail and sensorimotor cortical regions. This result is supported by previous rodent and human fMRI studies that showed that greater disengagement of the associative loop predicted higher learning performance ^8,49^ and proposed parallel but dissociable activity dynamics of the associative and sensorimotor loops. Furthermore, other modeling and fMRI studies have suggested a competitive interaction between the “fast” goal-directed and the “slow” habitual processes ^28,50-52^. Further studies would be still needed to elucidate the dynamic interactions between learning processes occurring at different—possibly multiple—time scales, as suggested by motor adaptation studies ^28,53^.

It should be noted that the current experiment provided feedbacks indicating current success (red-lighted target) or failure (unlighted target) without monetary reward for a successful performance. To provide monetary reward after a good performance has been shown to enhance skill consolidation ^54^ and even affect unconscious motivation ^55^, with heightened activation in the ventral regions of the basal ganglia. In a study which explored the dissociated effects of performance feedback and monetary reward on target-reaching performance using a force-grip device, feedback alone was sufficient to increase neural activity in the ventral striatum, while monetary reward was needed to elicit significant feedback-related increases in the activity of the dorsal striatum including the dorsal caudate nucleus ^56^. Interestingly, in the current study, we have observed robust success-modulated activities in the caudate nucleus in the absence of performance-dependent monetary reward. One possibility is that, in the case of a complex *de novo* learning paradigm which requires learning of an entirely new controller with considerable difficulty, positive feedbacks following successfully executed actions may be intrinsically considered more “rewarding” and would elicit strong neural activities in the caudate nucleus, especially in the head region. In contrast, when a relatively simple task involving more familiar or well-learned movements is presented, the caudate head may not respond as strongly to positive feedbacks, which would be associated with low intrinsic reward, unless the values of feedbacks are significantly heightened by the introduction of monetary reward.

There are several limitations of the current study. First, delineating the neural activity in the caudate tail using fMRI has been a challenge, due to its narrow and curved structure, proximity to the ventricles, and ensued partial volume effects ^57^. The current study attempted to bypass this issue at least partially by performing manual segmentation and using a high-resolution subcortical atlas. Nevertheless, the signals from the caudate tail are likely to be affected by partial volume effects and signals from nearby ventricles, which would in turn affect the current results at least to some degree. Thus, the current results regarding the caudate tail should be interpreted with caution. Yet, it should also be noted that evidence from animal studies, especially non-human primate studies, corroborates the presence of spatiotemporally distinct circuits in the caudate nucleus. Second, the current study does not provide an algorithmic explanation for possible mechanisms of *de novo* motor skill learning. While most model-based fMRI studies in reinforcement learning incorporated decision-making tasks with discrete state and choice spaces ^58-61^, the current study implemented a motor learning task that requires learners to make decisions continuously in high-dimensional state and choice spaces. Thus, learners were more likely to employ policy-based methods, directly mapping states to advantageous actions ^62,63^, instead of learning all the values of a combinatorially prohibitive number of state-action pairs in a continuous space. Finally, the current study also does not provide causal neurobiological accounts for the rostrocaudal transition in the caudate nucleus during motor skill learning. Future studies combining fMRI and noninvasive brain stimulation selectively targeting cognitive and sensorimotor networks may enhance our understanding of the intricate neural mechanisms underlying *de novo* motor skill learning.

## Materials and Methods

### Participants

Forty-three neurologically healthy young adults were enrolled in the current study. A total of 30 participants (12 females, mean age = 23.2 years; range = 19-30 years) completed all fMRI and behavioral experiment sessions and were included in the analyses. Among the 13 participants (3 females; mean age = 22.2 years; range = 18-28 years) who did not complete the entire experiment, three were excluded due to technical problems with data acquisition, and ten dropped out due to lightheadedness during fMRI sessions (n = 5), severe fatigue (n = 1), scheduling conflicts (n = 3), or failure to contact (n = 1). All participants were right-handed, according to a modified version of the Edinburgh Handedness Inventory ^64^. They had normal or corrected-to-normal vision and provided written informed consent approved by Sungkyunkwan University Institutional Review Board.

### Task procedure

We designed a novel task-based fMRI experiment for a complicated motor skill learning using an MR-compatible data-glove (14 Ultra, 5DT Technologies), based on a behavioral experiment implementing a data-glove paradigm ^65^. The data-glove measured finger flexures (two sensors per finger) and abductions between fingers (four sensors) from 14 sensors. The 14-dimensional vector (**h**) representing the hand posture, measured by the 14 sensors, was linearly mapped on to the two-dimensional position of the cursor (**p**) on the screen using the equation below ^23,44,62,65^

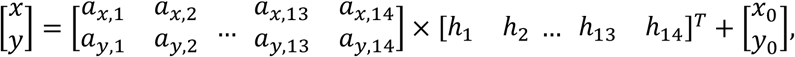

i.e., **P = Ah + P**_**0**_ where the mapping matrix **A** and the offset **P**_**0**_ were determined from the calibration phase in the first fMRI session. The time-series data were sampled from the 14 sensors at 60 Hz and each of the data was smoothed using the exponentially weighted moving average of 20 samples to reduce the intrinsic noise of the data-glove system. The smoothed data were used to construct the 14-dimensional vector **h**.

Stimuli were generated using a MATLAB toolbox Cogent 2000 (http://www.vislab.ucl.ac.uk/cogent.php) and were projected onto a custom-made viewing screen. Participants wore the data-glove on their right hand and placed the hand in a comfortable position, lying supine in the scanner and viewing the screen via a mirror. They were unable to see their hands throughout the experiment. Foam pads were applied to all participants to minimize head motions during the experiment.

### Experiment design

Participants completed two fMRI sessions and five behavioral training sessions between the fMRI sessions. The overall experiment typically lasted three weeks (range = 8-43 days, mean±SD = 23.4±8.7 days), and was completed within 45 days. The sessions for each participant were carefully scheduled so that the duration between the first behavioral training session and the second fMRI session would not exceed 15 days (range = 6-15 days, mean±SD = 9.9±2.7 days).

#### Visit 1

##### Familiarization and calibration

On the first visit, participants underwent a familiarization phase before fMRI scanning. Participants wore the data-glove on their right hands and were instructed to move their fingers freely for two minutes. A bar graph visualizing the real-time variance of each of 14 sensors was presented, to encourage participants to explore different movements. Then, principal component analysis (PCA) using the covariance matrix of the acquired time-series from 14 sensors was performed on the data. The first two principal components were used to construct the mapping matrix **A**, and the offset **P**_**0**_ was determined such that the mean hand posture mapped the center of the screen. Once the mapping was determined, the participants were instructed to reach all 25 targets of the 5×5 grid to ensure that all targets were reachable (Figure 1).

##### Resting-state fMRI scanning

One run of an 8-minute-long resting-state fMRI scan was acquired prior to task-based fMRI scans. Participants were instructed to keep their eyes open, maintain fixation on a cross presented in the middle of the screen, and refrain from focusing on any particular thought.

##### Localizer scanning

Localizer scanning was performed to define a region related to random finger movements. Participants were instructed to freely move their right fingers at natural speed when the text “Move” was presented and to stop when the text “Stop” was presented. Each “Move” or “Stop” condition lasted for one minute, and a total of four “Move”- “Stop” pairs were conducted. Using the finger movement data of the last two “Move” blocks, we recalibrated the mapping matrix **A** and the offset **P**_**0**_ as in the familiarization phase. We also confirmed that all 25 grid cells on the 5×5 grid could be reasonably reached by finger movements (Figure 1).

##### First main-task fMRI session

For each trial, a target appeared for 5 s as a gray grid cell with a yellow crosshair at its center in one of the four corners (1, top-left; 5, top-right; 21, bottom-left; 25, bottom-right) on the 5×5 grid. The cursor was displayed as a white crosshair. When the cursor reached the target grid, the color of the target grid changed to red. Holding a static posture led to the cursor staying at the approximately same location. The task was to place the cursor on the target grid as quickly and accurately as possible and maintain the cursor on the target. Participants were also instructed to move the cursor as straightly as possible when moving between targets.

There were seven runs in the first fMRI session, and each run consisted of 96 movements (composed of 8 blocks of 12 movements) from one target to another. The target sequence in each block was ordered as 1-5-25-21-1-25-5-21-25-1-21-5-1 and repeated for all eight blocks. In Runs 1-3, the mapping matrix **A** obtained earlier was implemented. In Runs 4-6, the two rows of the mapping matrix **A** were swapped such that the cursor position was flipped about the diagonal line. Then, in Run 7, the original mapping **A** was restored.

#### Visits 2-6

##### Behavioral training sessions

Participants performed five behavioral training sessions on separate days, each lasting about 40 minutes. The mapping implemented in the behavioral training sessions was the same as the initial mapping used in Runs 1, 2, 3, and 7 of the first fMRI session (i.e., “practiced mapping”). In each session composed of five runs, the first three runs presented the “main task,” which were identical to the task presented during the first fMRI session with targets appearing in the four corner cells of the grid. In the last two runs, participants were presented with the “sequential task,” in which they had to reach not just the four corner cells but also the cells consisting of the shortest path between each pair of the corner cells. For instance, participants had to reach Cells 1, 2, 3, 4, 5 consecutively, instead of simply reaching Cell 1 and then Cell 5 on the grid. Since the order of the corner-cell target sequence was the same as in the main-task blocks, the exact target sequence in each block was **1**-2-3-4-**5**-10-15-20-**25**-24-23-22-**21**-16-11-6-**1**-7-13-19-**25**-20-15-10-**5**-9-13-17-**21**-22-23-24-**25**-19-13-7-**1**-6-11-16-**21**-17-13-9-**5**-4-3-2-**1** (bold cell numbers indicating the corner cells). If the current target cell was reached for longer than 100 ms or not reached at all in 5000 ms, the target was changed to the next cell in the sequence. The goal of these additional runs was to encourage participants to move more straightly between targets.

#### Visit 7

##### Second main-task fMRI session

On the 7th visit, participants underwent the second fMRI session, which followed a procedure nearly identical to that of the first fMRI session. There were no familiarization and calibration phases. For the majority of participants, the second fMRI session was performed within 24 hours after the last behavioral training session. Otherwise, participants practiced the main task for a few minutes before the fMRI session.

### Behavioral data analysis

All statistical analyses and visualization were performed using Matlab (versions R2015b and R2018a, MathWorks, Natrick, USA), Python 3.6, and R 3.5.3. To obtain measures of learning performance, we calculated the success rates and aspect ratios of the cursor trajectories from two fMRI sessions and five behavioral training sessions.

#### Success rate

A trial-by-trial success rate was calculated as a proportion of time during the cursor was on the target, i.e., targets turned on in red. As each task block was designed to consist of all the 12 possible paths between four targets and the same target sequence was repetitive, we averaged the success rate in each block and estimated the learning rate (see Figure S1 for the details). In addition, we also calculated the overall success rate for each mapping, by averaging the block-by-block success rates from all fMRI and behavioral training sessions, and used it as individual participants’ learning performance.

#### Aspect ratio

For each of the trajectories between targets made by participants, we calculated its aspect ratio to find whether participants simply memorized the hand postures corresponding to targets or learned to control the cursor. Based on the description from the previous experiment using the data-glove paradigm ^65^, the aspect ratio was calculated by (i) estimating the maximum perpendicular distance between the straight line connecting two consecutive targets and the points on the actual cursor trajectory and (ii) normalizing the outcome with the length of the straight line.

#### Transfer of learning

The extent of transfer of learning between the practiced and unpracticed mappings was examined by the following procedure. First, as a measure of the amount of learning, the change in the success rate for each mapping in each fMRI session was obtained by subtracting the first-block success rate from the last-block success rate. The initial amount of transfer was then calculated by subtracting the learning amount for the practiced mapping in the first fMRI (“early-practiced”) from that for the unpracticed mapping in the first fMRI (“early-unpracticed”). The final amount of transfer was also calculated by subtracting the learning amount for the practiced mapping in the first fMRI (“early-practiced”) from that for the unpracticed mapping in the second fMRI (“late-unpracticed”). Finally, a Wilcoxon signed-rank test was performed to confirm whether the initial and final amounts of transfer were significantly different. We used a non-parametric test instead of a *t*-test due to the violation of normality identified in the initial transfer of learning.

#### Time-varying amount of movement

We calculated the time-varying amount of movement to examine whether it was collinear with the time-varying success rate used for the GLM analyses. For each of 14 sensors, we calculated the displacement (i.e., absolute change) for every 1 second and defined their average as the movement amount. For each of the three runs from the practiced mapping, we calculated Pearson’s correlation coefficients between the time-varying amount of movement and the success rate.

#### Duration of the experiment

The duration of the entire experiment in days, which might be considered as a potential confounding factor in the analyses of performance measures, did not show significant correlations with the overall success rate (*R* = 0.08, *p* = 0.665). It was thus not included as a covariate in the main analyses.

### 3-T MRI acquisition

We acquired fMRI data using a 3-T Siemens Magnetom Prisma scanner with a 64-channel head coil. Functional scans were acquired using an echo planar imaging (EPI) sequence with the following parameters: 1096 volumes (1113 volumes for resting state fMRI); repetition time (TR) = 460 ms; echo time (TE) = 27.20 ms; flip angle (FA) = 44°; field of view (FOV) = 220×220 mm; matrix, 82×82×56 voxels; 56 axial slices; slice thickness = 2.7mm. For anatomical reference, a whole-brain T1-weighted anatomical scan was performed using an MPRAGE sequence with the following parameters: TR = 2400 ms; TE = 2.34 ms; FA = 8°; FOV = 224×224 mm; matrix = 320×224×320 voxels; 224 axial slices; slice thickness = 0.7 mm. Before the functional scans, two EPI images with opposite phase encoding directions (posterior-to-anterior and anterior-to-posterior) were acquired for subsequent distortion correction, with the following parameters: TR = 7220 ms; TE = 73 ms; FA = 90°; FOV = 220×220 mm; matrix = 82×82×56; 56 axial slices; slice thickness 2.7 mm.

### fMRI data analysis

Analysis of fMRI data was performed using AFNI (Analysis of Functional NeuroImages, NIH, https://afni.nimh.nih.gov)^66^, Matlab (versions R2015b and R2018a, MathWorks, Natrick, USA), FreeSurfer (version 6.0.0., http://surfer.nmr.mgh.harvard.edu), Python 3.6, and R 3.5.3.

#### Preprocessing

Anatomical and functional image data were preprocessed using AFNI software (Analysis of Functional NeuroImages, NIH, https://afni.nimh.nih.gov)^66^. Task-based and resting-state functional images were first corrected for slice time acquisition and realigned to adjust for motion-related artifacts. Then retrospective distortion correction was performed using a field map calculated from the aforementioned two EPI images with opposite phase encoding directions. The corrected images were spatially registered to the anatomical data and transformed into Montreal Neurological Institute (MNI) template and resampled into 2.68 mm-cube voxels. All images were spatially smoothed through a Gaussian kernel of 4×4×4 mm full-width at half-maximum and scaled the time series to have a mean of 100 and range of 0 and 200.

#### Whole-brain voxel-wise GLM analysis

To identify regions responding to rewards, we designed a parametric regressor used in a subject-level general linear model (GLM) analysis (AFNI’s *3Ddeconvolve* function). Specifically, the reward amounts in the 1-second-long time bins, defined as the duration of the target grid cell reached successfully and turned red, were used as parameters modulating pulse regressors at the middle of time bins, and then convolved with a gamma function modeling a canonical hemodynamic response function (HRF). To better account for potential deviations from the HRF, a two-parameter Statistical Parametric Mapping gamma variate basis function with temporal derivatives (using *3Ddeconvolve*’s “SPMG2” option) was adopted for the main GLM results ^67^. For regressors of non-interest, we included six regressors estimating rigid-body head motion and five regressors for each run modeling up to fourth-order polynomial trends in the fMRI data. For each of the two fMRI sessions (“early” and “late” stages of learning), the GLM analyses were performed separately for practiced (fMRI runs 1-3) and unpracticed mappings (fMRI runs 4-6). Then, the regression coefficients of the parametric regressors were taken to a group-level whole-brain voxel-wise paired *t*-tests (AFNI’s *3dttest++* function) between conditions (early vs. late stages, practiced vs. unpracticed mappings). The voxel-wise threshold was *p* < 0.001, and the criterion of 40 suprathreshold voxels was determined by a conservative non-parametric method of randomization and permutation to provide a cluster-wise corrected threshold of *p* < 0.05 within the whole-brain group mask (AFNI’s *3dttest++* function with “-Clustsim” option). Notably, the results shown in Figure 3 and Table 1 were at much more stringent significance levels than the threshold ^68^.

#### Manual extraction of the caudate nucleus

To explore the distinct functional characteristics of the subregions of the caudate nucleus, we separated the caudate nucleus into three parts, head, body, and tail. Accurate delineation of the caudate tail is particularly difficult due to its narrow, curved, and obscured structure, which makes it inherently vulnerable to partial volume effects ^69^. Accordingly, the extremity of the caudate tail is frequently excluded from both atlas-defined and automatically segmented ROIs ^69^.

To better account for the problems regarding delineation as well as for individual variations in the caudate structure, we performed manual segmentation to create individual ROIs for the caudate nucleus. Manual segmentation for all participants was performed by one research trainee (S.Y.L.) using ITK-SNAP version 3.8.0 ^70^, with the guidance of researchers well experienced with MRI data processing and manual segmentation (H.F.K., H.J.K.). The boundaries of the caudate subregions were chosen based on previous literature ^57,69,71,72^. In each hemisphere, the caudate head ROI started from the region superior to the nucleus accumbens and extended along the anterior horn of the lateral ventricle, until it reached the vertical line traversing the anterior commissure (VAC) on the sagittal plane; the caudate body ROI, which continued to extend along the region superior to the body of the lateral ventricle, was bounded between the VAC and the vertical line traversing the posterior commissure (VPC) on the sagittal plane; the caudate tail ROI was defined as the region extending from the VPC back toward the anterior. To generate more even and continuous ROIs, additional postprocessing steps including small island removal (removing islands smaller than ten voxels) and smoothing (filling sharp corners and holes smaller than a kernel size of 3 mm) were performed using 3dSlicer version 4.10.2 (https://www.slicer.org/) ^73^.

As manual segmentation is also not free from the issue of partial volume effects, a separate set of caudate ROIs was created using the Reinforcement Learning Atlas ^74^, which provides high-resolution delineations of subcortical structures including the caudate tail. This set of ROIs was used for group analyses and visualization. The atlas-defined ROIs presented high similarity with the group average of manually segmented ROIs. Specifically, at the image intensity of 0.3 (an arbitrary threshold to maximize the similarity between the numbers of voxels contained by the manually segmented and corresponding atlas-defined ROIs), the group average of the combined manually segmented ROIs (663 voxels) and the combined atlas-defined ROI (662 voxels) demonstrated 61% overlap of selected voxels. All created ROIs were resampled using AFNI’s *3dresample* function to the participants’ EPI dimensions (2.683 mm^3^ isotropic voxels).

#### Region-of-interest (ROI) analysis

Based on the GLM results and previous literature on reward processing, the bilateral VMPFC, caudate head/body/tail, anterior/posterior putamen, globus pallidus (GP), and nucleus accumbens (NAc) were chosen as ROIs. The manually segmented caudate ROIs were used for the main analyses, and the VMPFC ROIs were extracted from the Automated Anatomical Labeling (AAL) 2 atlas ^75^. All other ROIs were generated using the Reinforcement Learning Atlas ^74^. The putamen ROIs were divided into the anterior (Y > – 0.5605) and posterior (Y < −3.2435) regions, with a 1-voxel gap between them to reduced partial volume effects, as suggested by previous literature ^76^.

For each ROI, the average beta estimates from the GLM analyses of success-rate modulation were extracted using AFNI’s *3dmaskave* function. The extracted data were then subjected to a two-way repeated-measures ANOVA, with the learning stage (early vs. late) and the mapping (practiced vs. unpracticed) as within-subject factors. For all such analyses, the effect sizes were estimated using partial eta-squared values, and subsequent post-hoc pairwise *t*-tests were performed with the Holm-Bonferroni adjustment to correct for multiple comparisons.

#### Resting-state functional connectivity analysis

After initial preprocessing, the resting-state fMRI data were further processed to control for white matter (WM) signals according to the following procedures. First, each participant’s white matter (WM) mask was created from automatic segmentation performed by FreeSurfer’s *recon-all* pipeline. Then, the resting-state fMRI time courses were detrended using fourth-order polynomial regressors and bandpass filtering (0.01-0.1 Hz) (AFNI’s *3dttest++* function). To avoid spurious correlation, we regressed out the first five principal components of signals from the WM mask ^77^. However, we did not perform global signal regression to avoid introducing artifactual anti-correlations ^78^. Importantly, signals from the ventricles were also not regressed out, in order to minimize the signal loss from the caudate tail, which is a region of interest located immediately next to the ventricles.

The resulting residual images were then used for the seed-based functional connectivity analysis. The seed ROIs were defined as the manually segmented individual ROIs of the caudate nucleus (bilateral head, body, and tail). We also independently defined three cortical ROIs: two ROIs in bilateral DLPFC for the rostral cognitive loop and one ROI in the left primary motor/somatosensory cortices (M1/S1) for the caudal sensorimotor loop. The DLPFC ROIs were defined as a mask obtained from Neurosynth, a large-scale meta-analytic fMRI database (http://neurosynth.org; accessed on September 23, 2019), with the use of the term “dorsolateral prefrontal,” which retrieved 1049 studies and 36216 activations. We then selected the two most significant clusters in the right and left prefrontal regions. For the ROI in the left M1/S1, we performed a whole-brain GLM analysis for the localizer fMRI data contrasting conditions between “Move” and “Stop” and then selected the most significant cluster in the left M1/S1 region (Figure S3B). The other six significant clusters related to finger movements are listed in Table S1.

For each participant, we calculated Pearson’s correlation coefficients between the mean time series extracted from each seed and residual signals from all other voxels in the cortical ROIs. The correlation coefficients were converted to Z-values using Fisher’s transformation (AFNI’s *3dTcorr1D* function with “-pearson -Fisher” options). Finally, for each cortical ROI, we concatenated the resulting Z-maps of the 29 participants (after excluding one outlier) and correlated them with individual learning performance measured as the overall success rate. We excluded one participant who had experienced technical issues that affected the performance during the behavioral training sessions and showed exceptionally low success rates during these sessions (lower 0.3 % in the distribution of performance; shown as the 29th participant in Figure S1).

## Data Availability

All MRI data used in this study are archived in the Sungkyunkwan University Network Attached Storage (NAS). The corresponding author can provide any information on the dataset related to this study if necessary.

## Acknowledgments

We thank Boohee Choi for assistance with data collection, Seung-Yeon Lee and Hye-Ji Kim for manual extraction of the caudate nucleus, and Hyung F Kim and Dongho Lee for their comments. Neuroimaging was performed at the Center for Neuroscience Imaging Research located in Sungkyunkwan University, supported by Institute for Basic Science.

## Author contributions

S.K. designed the study. Y.C., E.Y.S performed the experiment. Y.C., E.Y.S, and S.K. analyzed the data and discussed the results. Y.C. and S.K. wrote the manuscript.

