## Supplementary material for "Double dissociation of fMRI activity in the caudate nucleus supports *de novo* motor skill learning": Figure S1

**Figure S1. Individual learning performance for the practiced mapping across all sessions**

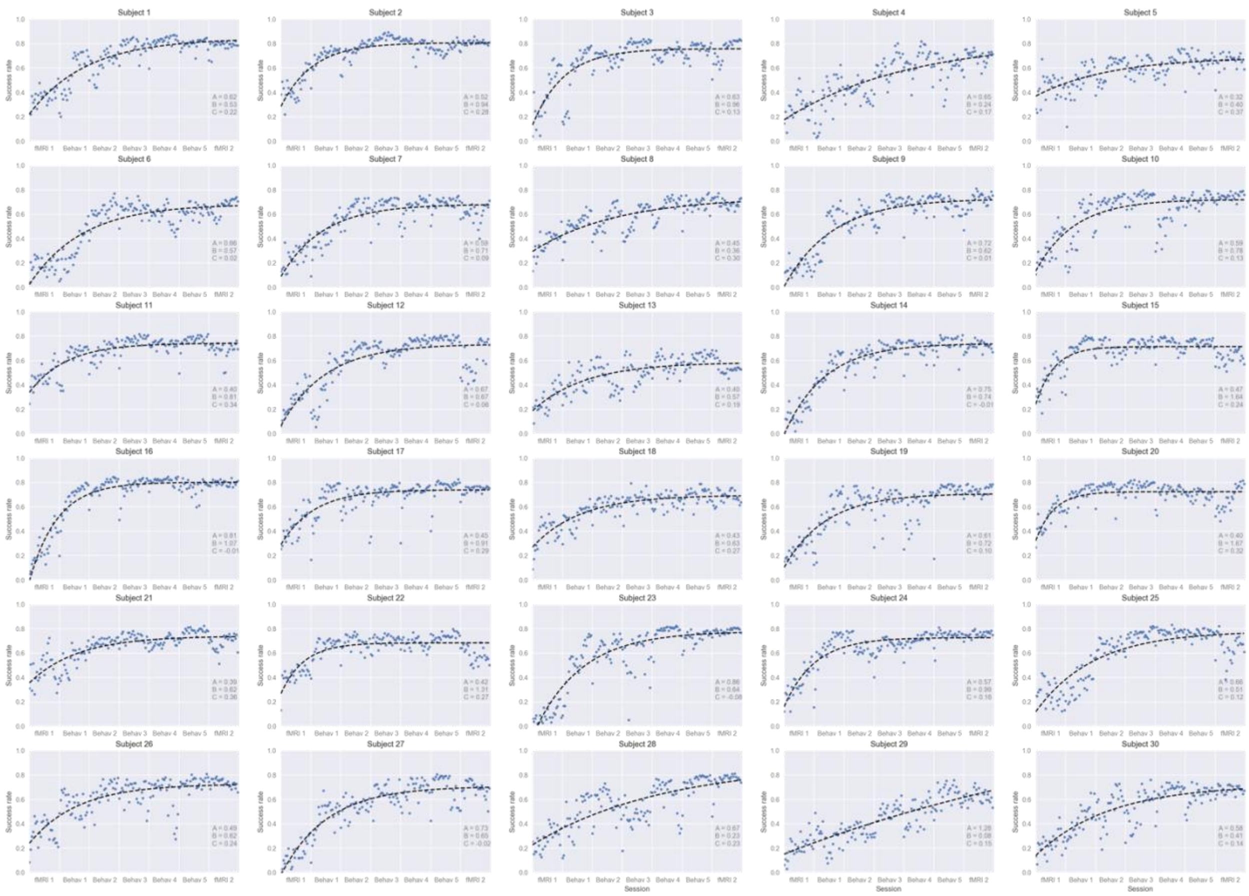

The individual success rate curves from all participants (n = 30). For the calculation of the individual learning rate, an exponential model with three parameters (*A*, *B*, *C*) was fitted to the block-by-block success rate for the practiced mapping for all fMRI and behavioral training sessions, with the following equation: $S(t) = A(1 - e^{-Bt}) + C$, where *t* denotes learning blocks, and *B* denotes the learning rate. Black solid lines denote the fitted exponential models. Blue dots indicate the actually observed block-by-block (consisting of 12 repetitive trials) success rates. The 29th participant was excluded as an outlier in the rs-fMRI functional connectivity analysis due to exceptionally low performance during the behavioral training sessions (lower 0.3% in the distribution of the overall success rates). The estimated learning rate of the outlier was also very low (0.076), compared with other participants (second lowest = 0.23). For each individual plot, the three parameters, A, B, and C, are shown in the bottom right.
