## Supplementary material for "Double dissociation of fMRI activity in the caudate nucleus supports *de novo* motor skill learning": Figure S2

**Figure S2. Whole-brain voxel-wise GLM analyses contrasting the early and late stages of learning (Late – Early)**

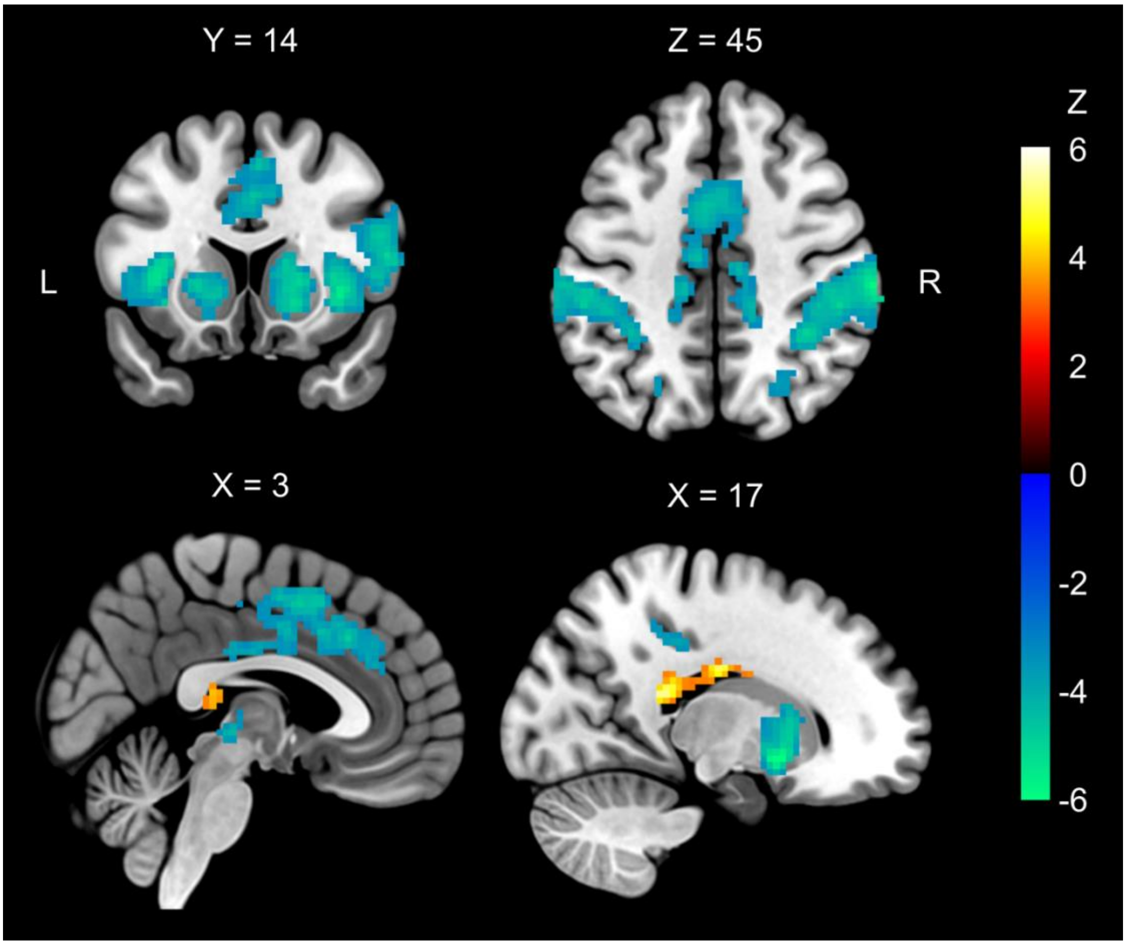


The regions colored from green to blue (including the bilateral supramarginal gyrus, middle cingulate cortex, insula, superior parietal cortex, rostral putamen, caudate nucleus) showed deceasing fMRI activities modulated by the success rates from the early to late stages of learning. In contrast, the bilateral caudate tail (colored from red to yellow) showed greater fMRI activities in the late stage. Color bars indicate the group-level *Z*-scores, and numbers above each slice indicate MNI coordinates.

Abbreviations: L, left; MNI, Montreal Neurological Institute; R, right.
