## Supplementary material for "Double dissociation of fMRI activity in the caudate nucleus supports *de novo* motor skill learning": Figure S3

**Figure S3. Locations of the independently defined ROIs encompassing the left/right DLPFC and left M1/S1**

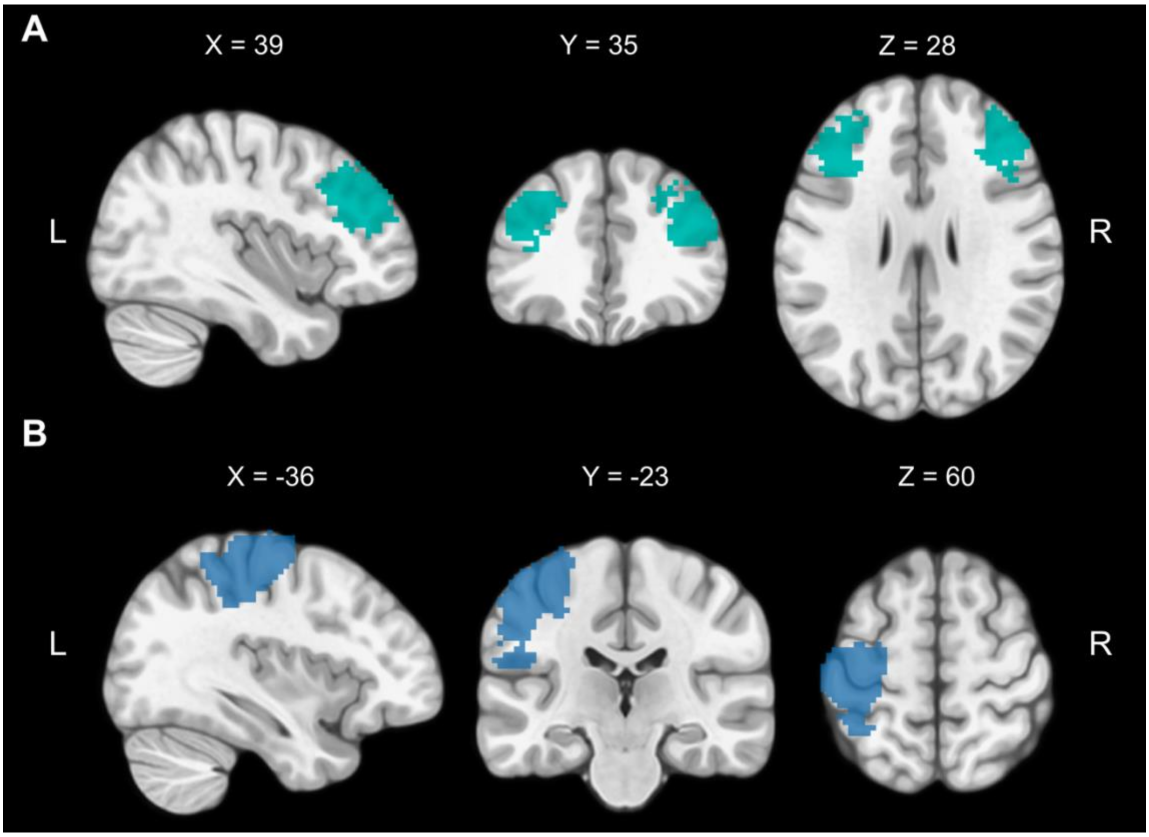


**A.** Cortical ROIs of the left/right DLFPC were obtained from the Neurosynth database (on September 23, 2019), with the use of the term “dorsolateral prefrontal” which retrieved 1049 studies and 36216 activations. **B.** The ROI encompassing the left M1/S1 was defined by using the data from an independent localizer scan, which identified the regions selectively activated by random movements of right fingers. The voxel activities were thresholded at a highly stringent level of significance, $p < 10^{-5}$. The XYZ coordinates are in the MNI space.

Abbreviations: DLPFC, dorsolateral prefrontal cortex; L, left; MNI, Montreal Neurological Institute; M1, primary motor cortex; R, right; S1, primary somatosensory cortex.
