## Supplementary material for "Double dissociation of fMRI activity in the caudate nucleus supports *de novo* motor skill learning": Table S1

**Table S1. Clusters showing activity significantly correlated with random finger movements**

| | Peak (MNI) | | | Cluster size (voxels) | *Z*-score at peak |
|---|---|---|---|---|---|
| | *X* | *Y* | *Z* | | |
| *L Pre/Postcentral Gyrus* | -42 | -23 | 60 | 1330 | 13.00 |
| *R Cerebellum, Lobules IV-V* | 15 | -53 | -26 | 500 | 13.00 |
| *R Postcentral Gyrus* | 47 | -26 | 46 | 338 | 6.22 |
| *L Posterior Putamen* | -31 | -7 | -2 | 331 | 6.44 |
| *R SMA* | 7 | -2 | 57 | 268 | 6.14 |
| *R Cerebellum, Lobule VIII* | 20 | -61 | -53 | 204 | 7.23 |
| *R Thalamus* | -15 | -23 | 3 | 175 | 6.51 |

Abbreviations: L, left; MNI, Montreal Neurological Institute; R, right; SMA, supplementary motor area.
